# A plant biodiversity effect resolved to a single genetic locus

**DOI:** 10.1101/264960

**Authors:** Samuel E. Wuest, Pascal A. Niklaus

## Abstract

Despite extensive evidence that biodiversity promotes plant community productivity, progress towards understanding the mechanistic basis of this effect remains slow, impeding the development of predictive ecological theory and agricultural applications. Here, we analysed non-additive interactions between genetically divergent Arabidopsis accessions in experimental plant communities. By combining methods from ecology and genetics, we identified a major effect locus at which allelic differences between individuals increases above-ground productivity of communities. In experiments with near-isogenic lines, we show that this diversity effect acts independently of other genomic regions and can be resolved to a single region representing less than 0.3% of the genome. Using plant-soil-feedback experiments, we also demonstrate that allelic diversity causes genotype-specific soil legacy responses in a consecutive growing period, even after the original community has disappeared. Our work thus suggests that positive diversity effects can be linked to single Mendelian factors, and that a range of complex community properties can have a simple cause. This may pave the way to a better understanding of diversity effects, and to novel breeding strategies, focussing on phenotypic properties that manifest themselves beyond isolated individuals, i.e. at a higher level of biological organisation.

---

More than two decades of plant ecological research and the publication of hundreds of studies have firmly established that positive biodiversity effects, in particular on community yield, are the rule rather than exception and are often substantial^1, 2^. These positive effects of biological diversity on community functioning have been explained by larger community-level resource use promoted by niche complementarity, facilitation, or by reduced negative density-dependent effects of enemies^3–5^. Yet our understanding of the particular driving mechanisms remains poor, for several reasons. First, diversity effects are emergent properties that only manifest in comparisons of communities differing in diversity^6^. Second, diversity effects, and the mechanisms that drive these, may change with environmental conditions^7, 8^. Third, while there is no doubt that functional trait differences underly biodiversity effects^9^, trait-based analyses remain to some degree phenomenological because evolutionary forces have led to the formation of trait syndromes, i.e. to sets of highly correlated traits that reflect fundamental trade-offs between ecological strategies^10, 11^. The observed variation in traits thus is confounded with evolutionary history^12^, and it remains almost impossible to distinguish the traits that are true drivers of biodiversity effects from traits that are merely correlated. It therefore remains difficult to develop predictive ecological theory and apply it, for example, in breeding and agriculture.

Most biodiversity research to date has focused on variation among species, but experimental^13^, theoretical^14^, and observational^15^ studies have shown that positive diversity effects on productivity also occur at levels of organization above (e.g. landscape level) and below species (e.g. genotype level). A substantial part of the trait variation apparent in plant communities occurs within species^16^, and increased intra-specific variation can have similar effects as inter-specific trait variation in low-diversity systems^13, 17, 18^. Despite qualitative differences, there may therefore be commonalities of trait variation within and between species with respect to effects and mechanisms, indicating that studies at the genotype level may provide some insights into effects of species-level variation and vice versa. A methodological advantage of intra-specific biodiversity studies is that genetic methods can circumvent some of the problems encountered in species-level diversity studies. Specifically, crosses between genotypes allow trait variation among individuals to be re-arranged^19^ without confounding with population structure or differentiation into ecological strategies. However, genetic approaches are normally used to study properties of individuals rather than emergent properties of communities. Here, we demonstrate in a case study how the genetic approach can be harnessed to identify the genetic underpinnings of biodiversity effects.

## Results

Using model plant communities (Fig. 1a), we screened ten pair-wise mixtures of genetically divergent natural accessions of *Arabidopsis thaliana* for which mapping populations are publicly available (Methods). Mixture communities of the two genetically divergent accessions Bayreuth (Bay) and Shadhara (Sha) reproducibly exhibited positive net biodiversity effects, i.e. mixtures produced a higher community-level shoot biomass than the average of their monocultures. This depended on soil conditions, with effects that were essentially absent on peat-rich soil but grew to an overyielding of 16% with increasing amounts of sand in the substrate (sand content × diversity: F_1,160_= 4.57, ANOVA P < 0.05; Fig. 1b and Supplementary Figure 1a). Analysis by additive partitioning^6^ revealed that these community-level biodiversity effects were due to complementarity rather than selection effects (sand content × complementarity effect, F_1,77_ = 7.21, ANOVA P < 0.01; Supplementary Figure 1b). Specifically, in our study both accessions benefited from growing in mixed communities.

**Figure 1.**
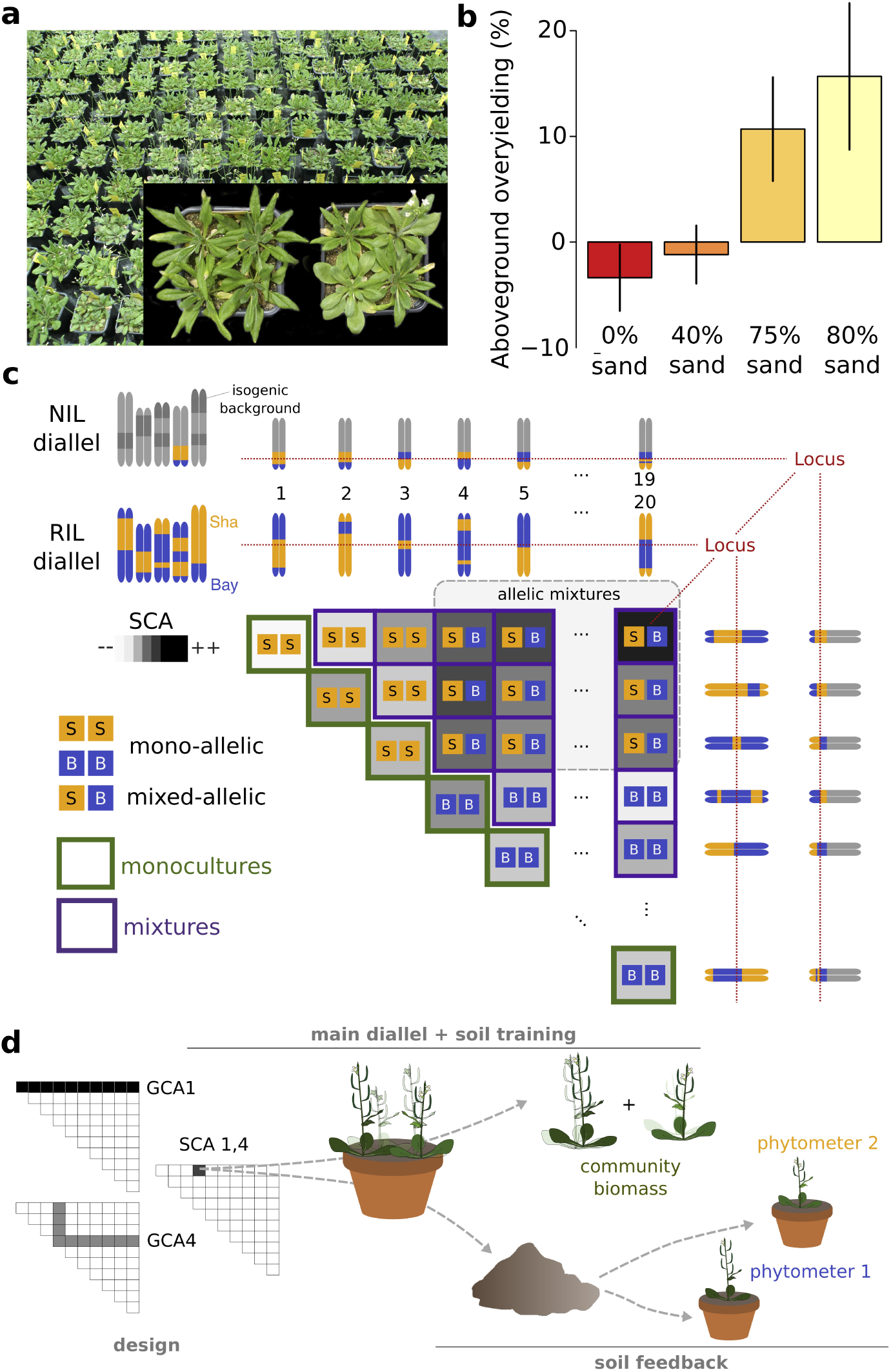
Combining ecological concepts and genetic methods. **a**, Pot systems used to study diversity effects in pair-wise genotype mixtures. The inset shows a Recombinant Inbred Line (RIL) monoculture (left) and mixture (right). **b**, Net diversity effects in Bay-Sha mixtures along a peat-sand substrate gradient. Error bars denote standard errors of means (s.e.m.). n = 164 pots **c**, Outline of the diallel design and the genotypes used throughout this paper. 18 RILs and the two parental accession, or 19 near-isogenic lines (NILs) were each placed in competition with each other, allowing to assess i) effects of genotypic mixture (i.e. diagonal vs off-diagonal), or ii) effects of allelic mixture at a given locus across all genotype mixtures (i.e. comparing SS and BB vs. SB) on pot productivity. **d**, Outline of the experimental procedure used in this work. Colored labels indicate measured variables. GCA = general combining ability; SCA = specific combining ability.

To analyse the genetic basis of the positive diversity effect in mixed Bay-Sha communities, we performed quantitative trait locus (QTL) mapping using publicly available recombinant inbred lines (RILs). These RILs are largely homozygous (Fig. 1c) and have been derived from a cross between the Bay and Sha accessions, followed by multiple subsequent rounds of selfing^20^. For efficient mapping, we capitalized on a so called competition diallel. Traditional diallel designs systematically cross sets of parental lines realizing all possible combinations and are used in breeding to determine the genetic basis of traits; specifically, diallel analysis partitions the trait variation of crosses into additive contributions^21^ of parental lines (general combining abilities; GCA) and cross-specific effects (specific combining abilities; SCA), with the latter interpreted as consequences of dominance or epistasis. By substituting individuals and crosses with communities and mixtures, the principle of diallels can be applied to the analyses of biodiversity effects in communities^22^, which we did here (Fig. 1c, d). In this context, the distinction between maternal and paternal effects ceases to apply, simplifying the design to a half-diallel. SCAs then quantify the deviations of mixture yields from expectations based on additive average contributions of the two genotypes. We combined 18 RILs and the two parental accessions Bay and Sha in a diallel, in four replicate blocks, on sand-rich soil. We detected significant positive genotype diversity effects on above-ground biomass production (Fig. 2a, F_1,189_= 10.47, P<0.01), indicating that the traits that promote biodiversity effects are heritable. As expected, a large proportion of the variation in SCA remained unexplained. We therefore tested for allelic diversity effects on SCAs at 69 marker positions. Both a marker regression technique and a standard QTL procedure revealed a major effect locus on the lower arm of chromosome four where allelic diversity at the community level resulted in higher SCAs (Fig. 2b and Supplementary Figure 2).

**Figure 2.**
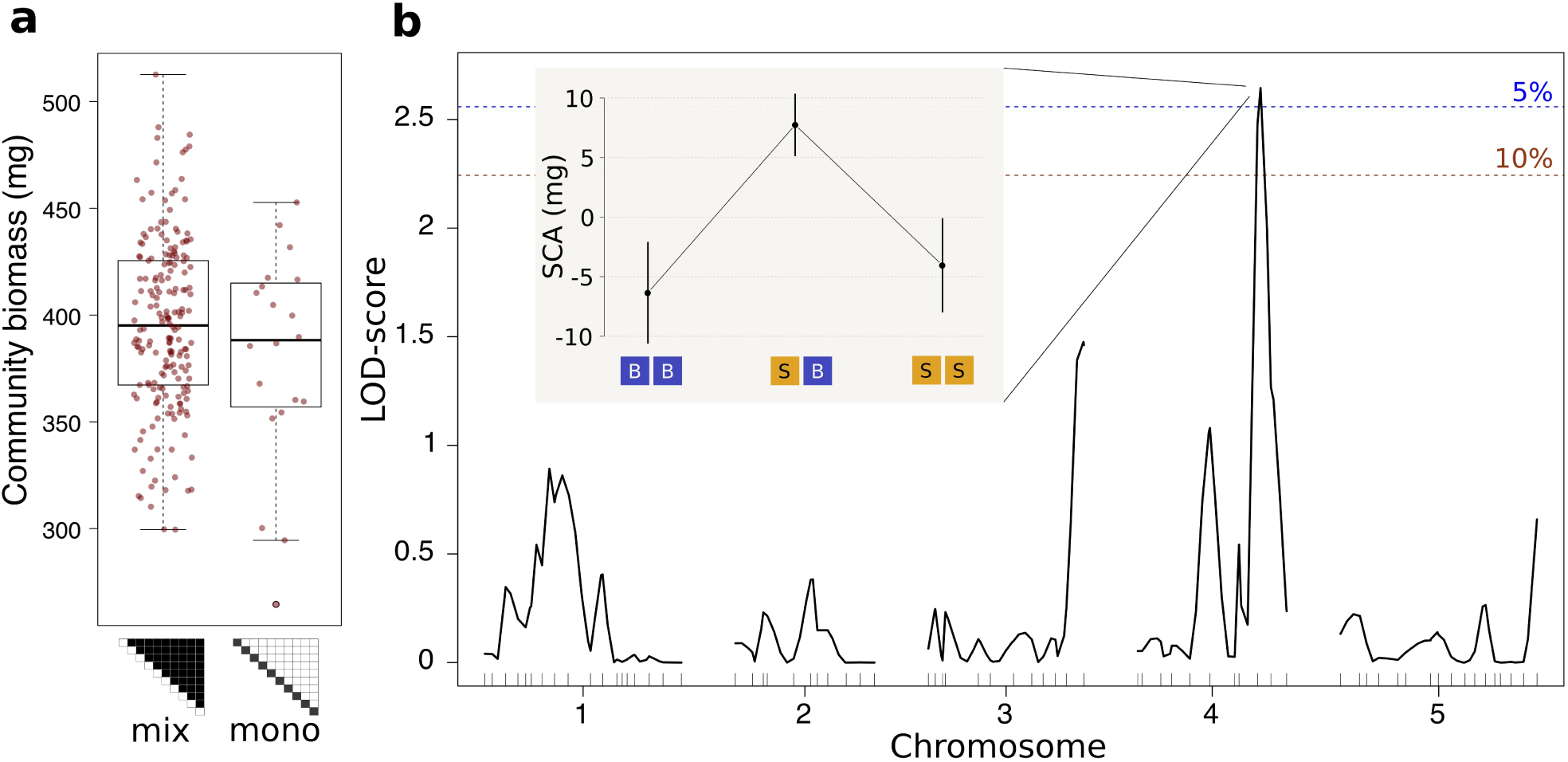
Allelic diversity at a major effect locus increases community productivity. **a**, Pot-level productivity in dependence of community type (mix = RIL mixtures vs mono = RIL monocultures), showing positive genotype mixture effects in the diallel and on sand-rich soil (values aggregated across four blocks, n = 871 pots / 210 compositions) **b**, Quantitative trait locus interval mapping of allelic diversity effects on specific combining ability (SCA). Vertical lines denote 10% and 5% genome-wide significance levels. The inset shows estimated SCA (± s.e.m.) across genotype mixtures that exhibit different allelic compositions at marker MSAT4.9 on chromosome four. LOD: logarithm of the odd.

With 18 recombinant lines, mapping resolution was limited and other effect loci or genetic interactions among loci may have gone unnoticed. We thus aimed at resolving the allelic diversity effect further to a single Mendelian factor. For this, we isolated a family of 19 near isogenic lines (NILs) that genetically varied only on the lower arm of chromosome 4, and in which we selected and inferred further recombination events by molecular markers and whole-genome re-sequencing (Fig. 3a,b, Supplementary Figure 3). With these NILs, we performed a second diallel experiment, replicated once on peat-rich soil where we expected no diversity effects and once on sand-rich soil where we expected positive diversity effects. Indeed, no locus was associated with positive allelic diversity effects on above-ground biomass on peat-rich soil (Fig. 3c). In contrast, on sand-rich soil we found a positive allelic diversity effects at a single locus (overyielding of 4.5%, Fig. 3d, P < 0.01), represented by a region of approx. 310 kb in size (termed locus Chr4@16.92: between 16.92 to 17.23 Mb). The overyielding of allelic mixtures of otherwise isogenic lines was transgressive: communities that contained individuals carrying different alleles at locus Chr4@16.92 in the NIL diallel produced more biomass than the most productive mono-allelic community (t = 2.32; P = 0.02), suggesting some form of functional complementarity between genotypes: relating to, without reference to a specific mechanisms at play, a phenomenon of two alleles positively interacting with each other when distributed *amongst* homozygous individuals of a community. Using structural equation models, we tested whether these allelic diversity effects were related to observed genotypic differences in shoot phenology or disease symptoms, but this was not the case (Supplementary Figure 4). Interestingly, however, this analysis showed that allelic diversity already manifested itself in increased community-level leaf cover early in the experient, when plants just started to produce flowering bolts (Supplementary Figure 4), indicating that the effect persisted through time. The allelic diversity effect we found, could, in principle, strictly depend on the genetic background. To rule out this possibility, we compared monocultures and mixtures of a second, independent pair of near-isogenic lines on both peat-rich or sand-rich soil (Supplementary Figure 5). Again, we found a significant allelic diversity effect on above-ground biomass that depended on soil (F_1,164_ = 4.17, ANOVA P=0.04 for soil x allelic diversity; 5.4% net overyielding on sand), and a significant effect of soil on the complementarity effect (F_1,82_=4.8, P=0.03). In conclusion, two consecutive diallel experiments using only 37 recombinant lines were sufficient to resolve a plant biodiversity effect to a genomic region representing ∼2.5‰ of the Arabidopsis genome (containing approx. 86 genes), which emphasizes the extreme efficiency of our approach. Our work suggests that biodiversity effects between genotypes can be dissected into discrete genetic elements that have major additive contributions.

**Figure 3.**
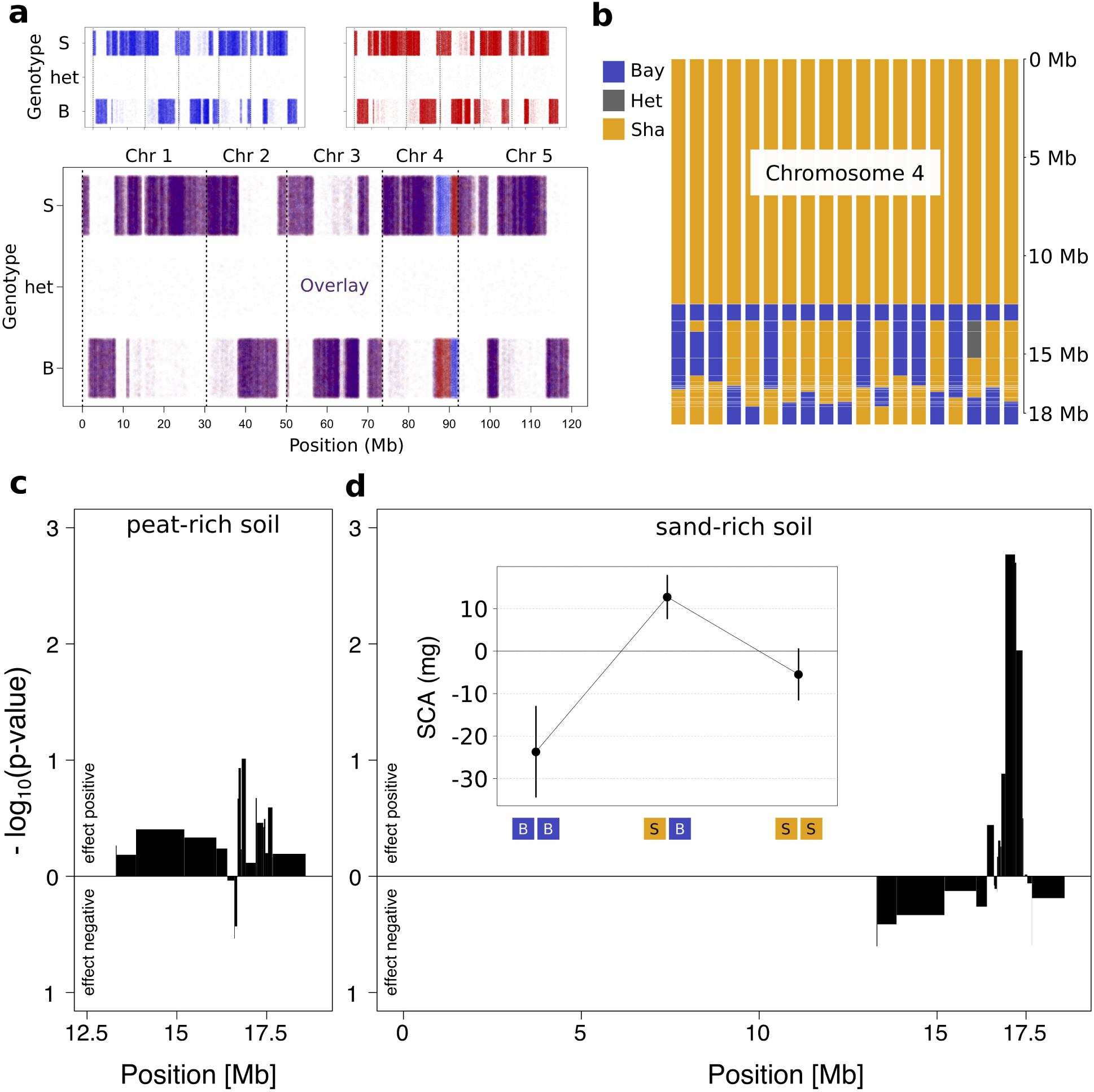
Resolving soil × allelic diversity interactions to a single Mendelian factor. **a**, Re-sequencing of near-isogenic lines (NILs) differing only on lower arm of chromosome four for fine-mapping. Shown are genotype calls at all polymorphic sites across the genome (B = homozygous for the Bay allele, het = heterozygous, S = Sha allele) in either NIL r10 (blue, top left) or NIL r96 (red, top right), as well as an overlay of the two line’s genotype calls (bottom). **b**, Reconstructed genotypes across chromosome four of the 19 NILs used for fine-mapping. Each bar represents a single NIL. **c, d**, Map of allelic diversity effects across chromosome four, on either peat-rich soil (c), or sand-rich soil (d). The widths of the bars indicate the size of the regions in which no recombination events were inferred across the whole population. Inset in (d) shows the mean ± s.e.m. of SCAs across allelic compositions at the diversity effect QTL.

**Figure 4.**
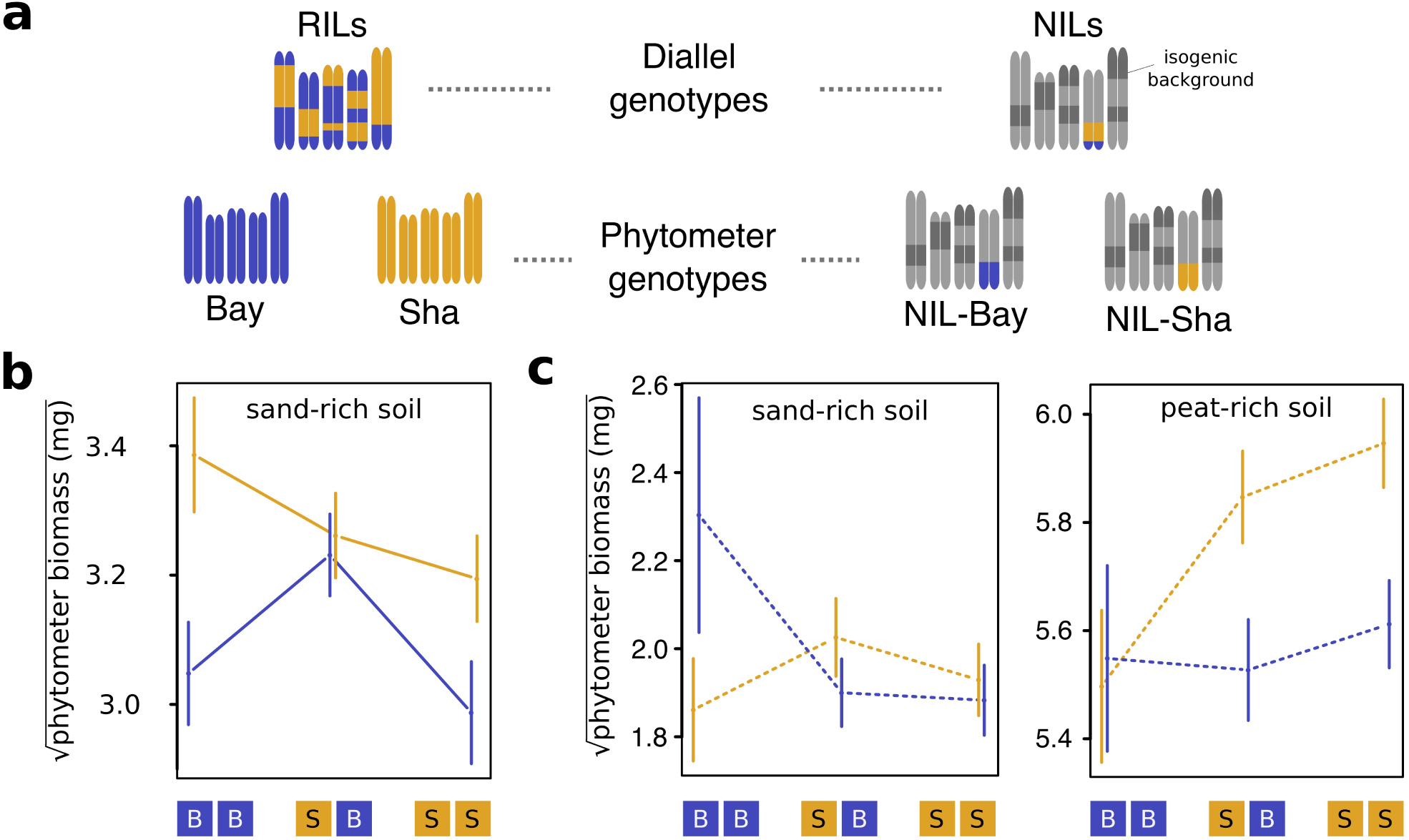
Allelic diversity effects persist across a generation through their soil legacy. **a**, Scheme of the genotype-combinations used for either the recombinant inbred lines (RILs) vs. near-isogenic lines (NILs) diallels (top), and the phytometer genotypes used in the soil-feedback phase (bottom). **b**, phytometer performance (Bay or Sha, mean ± s.e.m.) on legacy soil derived from RIL mixtures with different allelic compositions at marker MSAT4.9. **c**, phytometer performance (NIL-Bay or NIL-Sha, mean ±s.e.m.) on legacy soil derived from NIL mixtures with different allelic compositions at locus Chr4@16.92 on either sand-rich (left) or peat-rich (right) soil. Values were square-root transformed for analyses, n = 851 and 720 pots for RIL and NIL diallels, respectively.

There is growing evidence that productivity responses in plant biodiversity experiments^4, 5, 23, 24^ and their increase through time^25^ are related to diversity-dependent soil conditioning. To test whether allelic diversity causes effects through soil conditioning also in our study, we performed soil feedback experiments. Our objectives were twofold: first, we were interested whether there were general effects of allelic diversity on soil quality; second, we aimed to test whether allelic interactions occurred among plants that did not grow simultaneously, i.e. whether allelic effects were mediated through time by soil legacy. We assessed soil conditioning by growing indicator plants (“phytometers”^25^) on soil collected from both diallel experiments^26^ when these were harvested (Fig. 1d and Supplementary Discussion). The phytometers were the two parental accessions Bay or Sha for the RIL diallel, and two near-isogenic lines in the NIL diallel.

We indeed found phytometer-specific soil legacy responses that depended on the allelic diversity of the communities that had conditioned the soils in the preceeding growing period (Fig. 4; RIL diallel: diversity at marker MSAT4.9 × phytometer; F_1,166_= 6.48; ANOVA P = 0.012; NIL diallel: diversity at locus Chr4@16.92 × phytometer; F_1,168_= 5.61; P = 0.02; Supplementary Table 1; Supplementary Figure 6). These phytometer-specific responses to soil legacy were independent of differences in previous community productivity and associated resource depletion (the effects remained statistically significant and comparable in size when first adjusting for community biomass in linear models). However, the effects differed between phytometers and experiments, i.e. they depended on environmental or genetic context (Supplementary Discussion). This is not surprising in light of the complex mechanisms involved. Developing a full understanding of the biological mechanisms at play will thus require further experiments, including soil analyses. Nonetheless, these experiments demonstrate that allelic differences at a single QTL cause interactions between individuals within a community and also, mediated by a soil-borne factor, through time. The latter can be perceived as an “extended phenotype” sensu Dawkins^27^, the expression of which depends on interactions between group members. We were intrigued to find that we could, in principle, have genetically mapped the allelic diversity effect solely through its soil legacy; in other words, by QTL mapping this extended phenotypic property of allelic mixtures (Supplementary Figure 6e,f; Supplementary Discussion).

## Discussion

Our study systematically resolves a biodiversity effect, once identified in a specific set of interacting plants and environmental context, to between-individual allelic differences in a single chromsomal region So far, complex emergent properties of plant communities did not necessarily seem genetically tractable, especially since quantitative traits of individuals often are polygenic^28^ if not omnigenic in nature^29^. A single case study obviously is limited with respect to generalizations, but we consider it possible that in many cases between-individual allelic complementarity and resulting biodiversity effects might instead have a relatively simple genetic architecture – a feature not uncommon for other types of biotic interactions^30^. Our genetic approach is extensible to the study of interactions among other genotype combinations, and, with modifications, among species, and could thus lead to fundamental new insights into the traits and genetic underpinnings of biodiversity effects in more natural systems. Equally importantly, the genetic tractability of such effects may allow efficient breeding of genotype mixtures that support increased yields through some form of functional complementarity while maintaining low variation in economically relevant traits. Biodiversity effects have received relatively little attention in breeding and conventional agriculture, with the notable exception of crop rotation^31^ and intercropping of cultivars and species^32–34^. Instead, sustaining a growing global human population heavily depends on increasing nutrient inputs to crop production systems^35^, on breeding of single genotypes for monoculture performance^36^, and on the use of within-individual diversity effects termed heterosis^37^. Our approach might help bypass constraints imposed on the performance of single genotypes, by shifting breeding efforts from the individual to the system level^38^.

## Methods

### Germplasm

The Shadhara and Bayreuth accessions were kindly provided by Nuno Pires (University of Zurich) and had originally been obtained from the Nottingham Arabidopsis stock center (NASC). We selected these two genotypes following an initial screen of ten pairs of accessions, because they exhibited an above-average but not extreme biodiversity effect when growh together on sand-rich soil, and because a high-quality and frequently used RIL population is publicly available^20^. It also was interesting that the biodiversity effect vanished on peat-rich soil. The 18 RILs (representing the “RIL-minimal set” and line 33RV191 used to generate NILs (all contained in the core-pop set of 165 lines) were ordered from the Versailles Arabidopsis stock center (*http://publiclines.versailles.inra.fr*) and propagated in a growth chamber. A Bay×Sha RIL (33RV191) was confirmed to be heterozygous at two PCR marker positions on chromosome 4 (Table S2). Upon selfing of this line, the two NILs 33RV191-Sha and 33RV191-Bay were isolated (referred to as NIL-Bay or NIL-Sha hereafter), and their genomes were re-sequenced as described below. Using the same procedure, an independent pair of near-isogenic lines was derived upon selfing the Bay×Sha RIL 33RV77, giving rise to NILs 33RV77-Sha and 33RV77-Bay. Furthermore, after selfing of another single heterozygous F_10_ individual of line 33RV191, we screened 160 offspring for recombination between the ShaBa5, ShaBa6 and ShaBa8 markers on chromosome four. Upon selfing of 23 putative recombinant offspring, we isolated 19 homozygous recombinant lines for which we confirmed a recombination event in the region by PCR. We then performed whole-genome re-sequencing to confirm the isogenic background and to infer recombination breakpoints for this heterogeneous inbred family (referred to as NILs throughout the text) as described below.

### Soils and growth conditions

Soils consisted of different mixtures of a peat and nutrient rich soil (Einheitserde ED73; pH ∼5.8, N 250 mg L^-1^; P_2_O_5_300 mg L^-1^; 75% organic matter content; Gebrüder Patzer GmbH, Sinntal-Jossa, Germany) and finely grained quartz sand. Pot for all mixture experiments were 7×7×8 cm in size. The experiment using the parental lines Bay and Sha was replicated on a soil quality gradient with sand contents of 0%, 40%, 75% and 80%, which resulted in a near-linear decrease of pot-productivity from the highest to lowest ED73 content, likely through a dilution of soil nutrients (Fig. S1). For the rough-mapping of the diversity effect using RILs, we used a mixture of 80% sand and 20% ED73. For the fine-mapping diallel using NILs, we used either a 80%:20% or a 20%:80% sand:ED73 mixture.

Seeds were sown directly onto soils (approx. 10 seeds per position, 4 positions per pot, Fig. 1a). The pots were placed in growth chambers or greenhouse compartments and covered with plastic lids to maintain a high humidity for germination and initial seedling establishment. Additional light was provided if necessery, achieving a photoperiod of 14–16 hours. Day-time and night-time temperatures were maintained around 20–25 °C and 16–20 °C, respectively. Seedlings were thinned continuously until a single healthy seedling remained per position.

Once seedlings were established, the pots were placed in a greenhouse compartment with automated watering (every 2 days). In summer 2015, daytime temperatures were extremely high, and the first block of the RIL diallel was therefore grown in a growth chamber with full climate control (8 h night/16 h day; 60% humidity; 18/23°C night/day temperature). The second block was grown in the growth chamber for a month before it was re-located to the regular greenhouse compartment. Pots that did not contain all four originally planted individuals were discarded. Plants were harvested 43–51 days after sowing, with the specific harvest date determined by the occurence of approx. 5–10 dehiscent siliques on the earliest flowering genotypes within a block.

After the diallels were harvested, soil feedback trials were established by dividing the soil of a pot into two smaller pots 5.5×5.5×6.0 cm in size. The respective phytometers (Bay or Sha for the RIL diallel, 33RV191-Sha or 33RV191-Bay for the NIL diallel) were sown directly onto the soil. Again, seeds were oversown and seedlings thinned continously until a single healthy individual remained. Phytometer experiments were harvested either 36 days after sowing (peat-rich soil remaining after the NIL diallel, harvested early because plant roots started to grow out of the pots) or 49–58 days after sowing (sand-rich soil, each block was harvested on a single day). For all experiments, the position of the individual pots was randomized across trays during seedling establishment, and across watering tables after seedling establishment. Throughout the experiment, pots were re-positioned randomly within trays and tables every 7–10 days. Pots were watered *ad libitum*, and in case of high population densities of dark-winged fungus gnats, the systemic insectizide ActaraG (Syngenta Agro AG) was applied according to the manufacturers recommendation. After harvesting, plant biomass was dried at 65°C for at least three days before weighing. We determined early rosette cover in the NIL diallel by photographing pots 27 days after sowing and estimating the horizontally projected community-level rosette area using the Easy Leaf Area software^39^. We further recorded the occurrence of leaf disease symptoms (wilting, blotching, or early senescence) 30 days after sowing. As a proxy for flowering time in the NIL diallel, we measured flowering bolt height of all plants 35 days after sowing; by then, >98% of the individuals had a flowering bolt longer than 0.5 cm). The NIL diallel was harvested 50 days after sowing.

### Experimental designs

To test the soil-dependency of biodiversity effects in mixed Shadhara-Bayreuth communities, four soil substrates varying in sand content were prepared as described above. We then grew 12 replicate monocultures of each accession plus 24 replicate mixtures per soil type (total of 48×4= 192 pots). The RIL diallel consisted of a half diallel replicated in four blocks. All pair-wise RIL combinations were realized once per block except for RIL monocultures which were replicated twice. For the follow-up soil feedback experiment, we re-used soil from only the first two blocks of the diallel. We re-mixed the soil of each single pot after harvesting the plants, and re-distributed it into two smaller plots that were sown with either a Shadhara or Bayreuth parental genotype that served as phytometers.

The NIL diallel used for fine-mapping was realized in a single block that contained all pair-wise combinations of the 19 NILs including monocultures. The subsequent soil feedback part of the experiment was realized as described for the RIL diallel, using either 33RV191-Bay or 33RV191-Sha genotypes as phytometers. To test if the allelic diversity effect was strictly dependent on genetic background, we grew 21 replicate monocultures of each NIL 33RV77-Bay or 33RV77-Sha, plus 42 replicate mixtures, on either peat-rich (80% ED73, 20% sand) or sand-rich (20% ED73, 80% sand) soil (total of 42×2×2 = 168 pots).

### Genotyping and line re-sequencing

PCR-based genotyping assays (Table S2) were developed based on deletions in the Sha genome as predicted by the Polymorph tool (http://polymorph.weigelworld.org)^40^.

Barcoded libraries for genome re-sequencing were prepared using the Illumina Nextera DNA Library Prep Kit (FC-121-1031, Illumina Inc. San Diego, CA) in combination with the Nextera Index Kit (96 indices, FC-121-1012) and pair-end sequenced on an Illumina HiSeq 2500 (2×150 bp, rapid run). The clustering and sequencing were performed at the Functional Genomics Center Zurich. Sequences were aligned to the Arabidopsis genome (Col-0 genome, TAIR version 10) using BWA^41^, aligned read sorting and variant calling were performed using samtools^42^. Aligned genomic sequences of the parental accessions Bay-0 and Sha were downloaded from the 1001 genomes project data center (http://1001genomes.org). The VCF-file produced by the samtools software was loaded into the R Statistical Software^43^, where the subsequent analyses were performed: variant calls were filtered (for differences in genotype calls between the Sha and Bay genomes, quality of variant calls, population-level minimal minor allele frequency 0.2; maximum heterozygosity 0.2). Inference of genotype calls at polymorphic sites was performed as described previously^44^ and inference of parental alleles was improved using functionality implemented in the MPR package^44^. Genotype reconstruction was then performed in R using a simple hidden Markov model as implemented in the R package HMM, with hidden state starting probabilities (Bay, Het or Sha) all set to 1/3, and transition probabilities from one state to itself set to 0.99998 and to the other two states set to 0.00001 each. Emission probabilities of genotype calls given a state, e.g. Bay, were set to 0.35, 0.25, 0.25, 0.15 for genotypes calls Bay, Het, Sha or missing, etc.

### Statistical analyses

We analyzed data from the diallel experiments using linear mixed models summarized by analysis of variance (ANOVA). The model terms included, in this order, the general combining abilities (GCA) of genotypes (a factor with 20 levels in the RIL diallel and 19 levels in the NIL diallel), the genotype diversity in the pot (GD, 1 or 2 genotypes), the allele identity in the genotype monocultures (A, Sha or Bay), the allelic diversity in the genotype mixtures (AD, 1 [Sha/Sha or Bay/Bay] or 2 [Sha/Bay]), and the genotype composition planted in the pot (comp). The factor GCA was created by superimposing the model matrices for factors coding for the first and second genotype (factors with 20 and 19 levels for RIL and NIL diallels, respectively). The significance of GD, A, and AD were determined using F-tests with comp as error term (denominator). A and AD were encoded in such a way that these contrasts applied only to genotype monocultures and mixtures, respectively. Technically, this was achieved by including a third level in the factor that did not vary in the other group. Fitting A and AD after GD therefore only explained variance in these subsets. The diallel model was extended by additional terms and the corresponding interactions when these applied; specifically, the RIL diallel included a block effect. The NIL diallel included terms for soil type, and interactions of all the terms above with soil type (for example, soil×AD was tested using soil×comp as error term). The soil feedback experiments included further interactions with phytometer (RIL and NIL diallel), and phytometer×soil (NIL diallel). Effects of pot biomass in the diallel (diallel biomass and diallel biomass × soil) were accounted for in these linear models, and data were square-root transformed to obtain normally distributed residuals.

Specific combining abilities for mapping were calculated directly, within blocks, by solving the linear model m = X GCA + SCA where X is the design matrix describing the genotype composition of a pot. Monoculture SCAs were also determined but not used for QTL mapping of allelic diversity within RIL mixtures. In the RIL diallel, the SCAs of each genotype composition was first calculated per block and then aggregated over all blocks using least-square estimates. Marker regression was performed contrasting SCAs of mono-allelic RIL mixtures (“BB” and “SS” compositions) with mixed-allelic mixtures (“BS” compositions) using the glht-function provided by the multcomp package^43^. QTL mapping was also performed using the R/qtl package and interval mapping (scanone-function), with both mono-allelic compositions at a given locus re-coded as to the same level (“mono-allelic”) and compared against mixed-allelic compositions. Genome-wide significance was assessed by resampling (n=5000).

To test the relationships of allelic diversity effects on SCAs with measured traits, we developed multip-group (sand-rich and peat-rich soil) structural equation models using lavaan (http://lavaan.ugent.be). Allelic diversity at Chr4@16.92 was included as exogenous variable; endogeneous variables were two metrics of trait variation among genotypes (which are possible indicators of complementarity), and early community-level projected leaf area (see above). Trait variation was quantified as difference among bolt length of the genotypes (square-root-transformed), and as difference in the occurence of leaf disease symptoms (a binary variable). Starting with a near-saturated model, the modelled paths were simplified in an educated way until a minimal model was found for which the model-implied and observed covariance structure among variables did not differ significantly (X^2^-test).

## Acknowledgements

We thank Bernhard Schmid and Ueli Grossniklaus for helpful discussions and sharing infrastructure. We thank Jordi Bacompte and Jacob Weiner for helpful comments on the manuscript. We further acknowledge Matthias Philipp for technical support, Enrica De Luca and Nicole Ponta for help with plant measurements, Matthias Furler and Dorde Topalovic for technical greenhouse support. This work was supported by an Ambizione Fellowship (PZ00P3_148223) of the Swiss National Science Foundation (to S.E.W). P.A.N acknowledges support by the University of Zurich Priority Programme “Global Change and Biodiversity”. S.E.W was also financially supported by funds of the University of Zurich and the European Research Council (to Ueli Grossniklaus).

## Author contributions

S.E.W. conceptualized and designed the research (with input from P.A.N.) and performed the experiments. Both authors performed the analyses and wrote the manuscript. Both authors revised and approved the final version of the manuscript.

## Data availability

The datasets described in the paper and a functional annotation of the 86 genes within the fine-mapped diversity QTL are available through the Zenodo data repository (DOI:10.5281/zenodo.1254563). Sequencing data are deposited in the NCBI Short-Read Archive (accession SRP149077).

Analysis scripts are available from the authors upon request.

## Supplementary Tables

**Supplementary Table 1.**
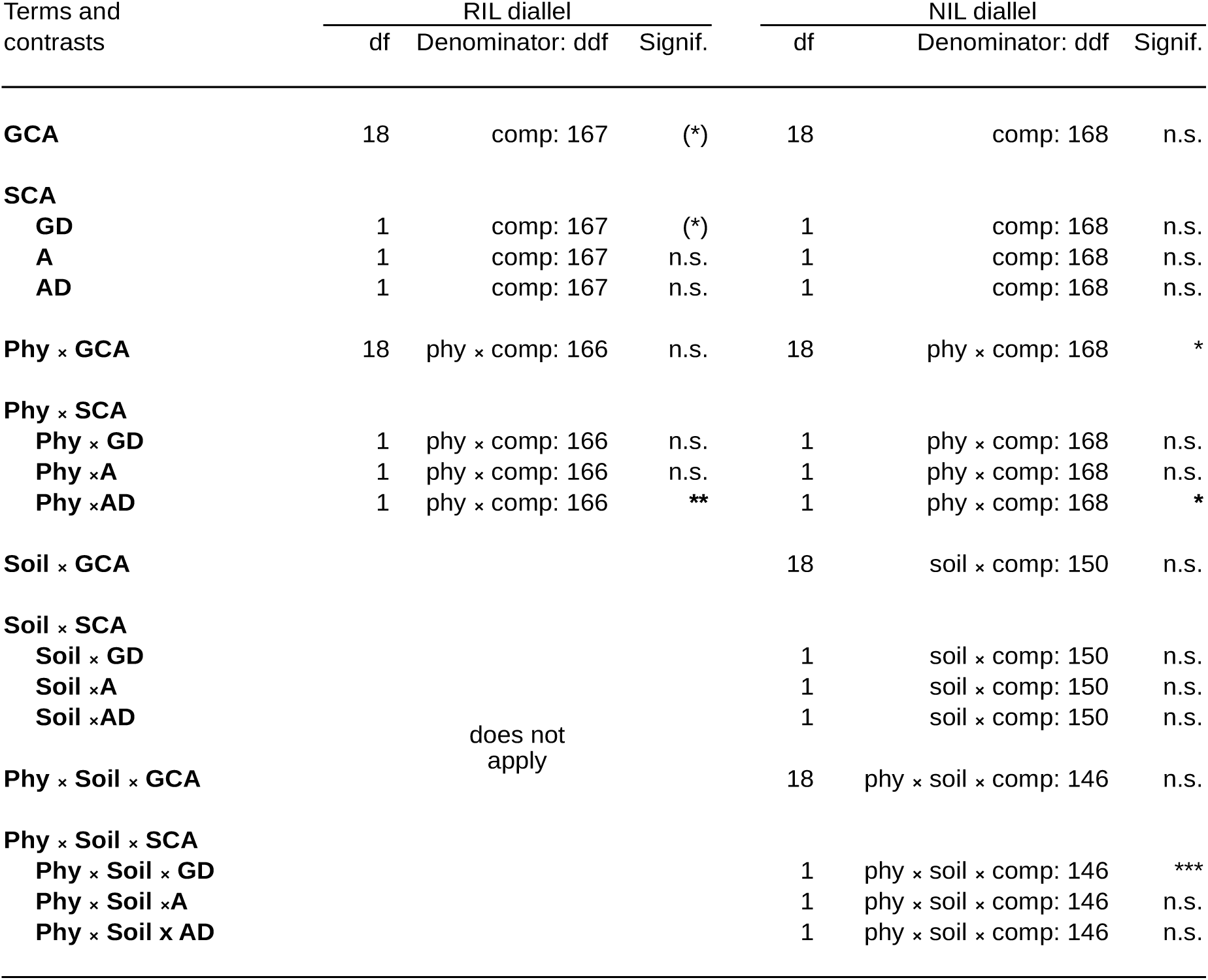
Legacy effects of soils conditioned in RIL and NIL diallel experiments. Effects were quantified using two phytometers (factor ‘phy’; Sha and Bay accessions in RIL diallel, and near-isogenic lines bearing Sha and Bay allele at putative effect locus in NIL diallel). The NIL diallel additionally was replicated on substrates differing in sand content (factor ‘soil’). The term ‘GCA’ (general combining abilities) indicates average genotype-specific soil conditioning effects on phytometer yields. ‘SCA’ (specific combining abilities) captures deviations in yield from additive predictions made using GCAs. Within SCA, the following contrasts were tested: GD: Genotype diversity, i.e. whether genotype monocultures differed in feedback effects from two-genotype mixtures; A: allele-specific differences at marker MSAT4.9 (RIL diallel) and Chr4@16.92 (NIL diallel) within genotype monocultures; AD: allele-diversity effects within genotype mixtures. df and ddf indicate nominator and denominator degrees of freedom of corresponding F-tests. *** P<0.001; ** P<0.01; * P<0.05; (*) P<0.1; n.s. not significant.

**Supplementary Table 2.**
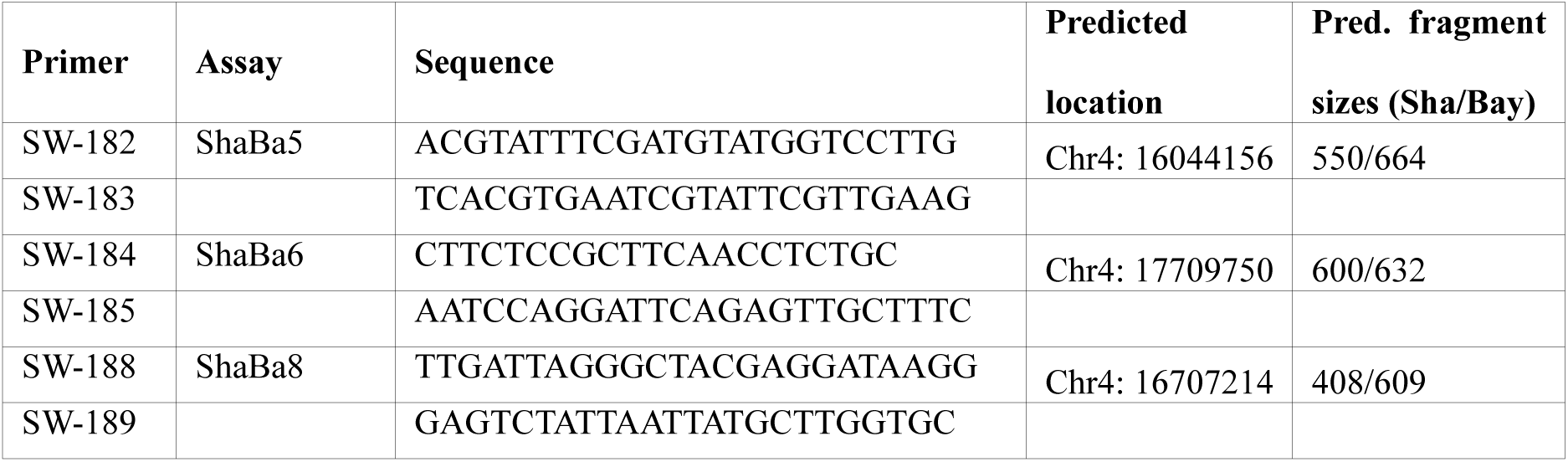
PCR markers used in this study.

## Supplementary Figures

**Supplementary Figure 1.**
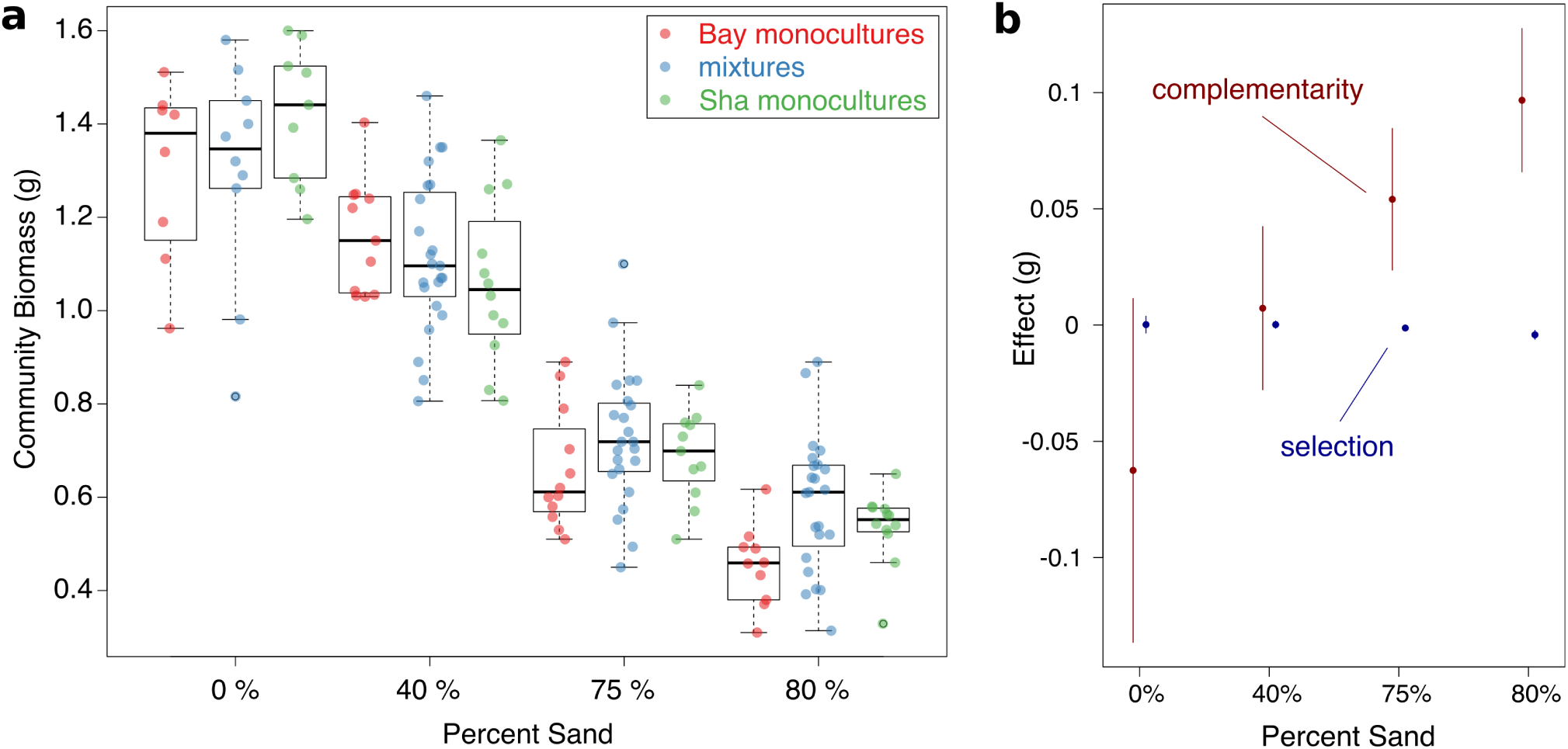
Productivity and complementarity in Bay-Sha mixtures across a peat-sand gradient. **a**, Pot-level biomass measurements of Sha and Bay monocultures or pair-wise mixtures along a sand/peat substrate gradient. n = 164 pots **b**, Complementarity and selection effects calculated according to the additive partitioning ^1^ method along the substrate gradient. Error bars denote s.e.m.

**Supplementary Figure 2.**
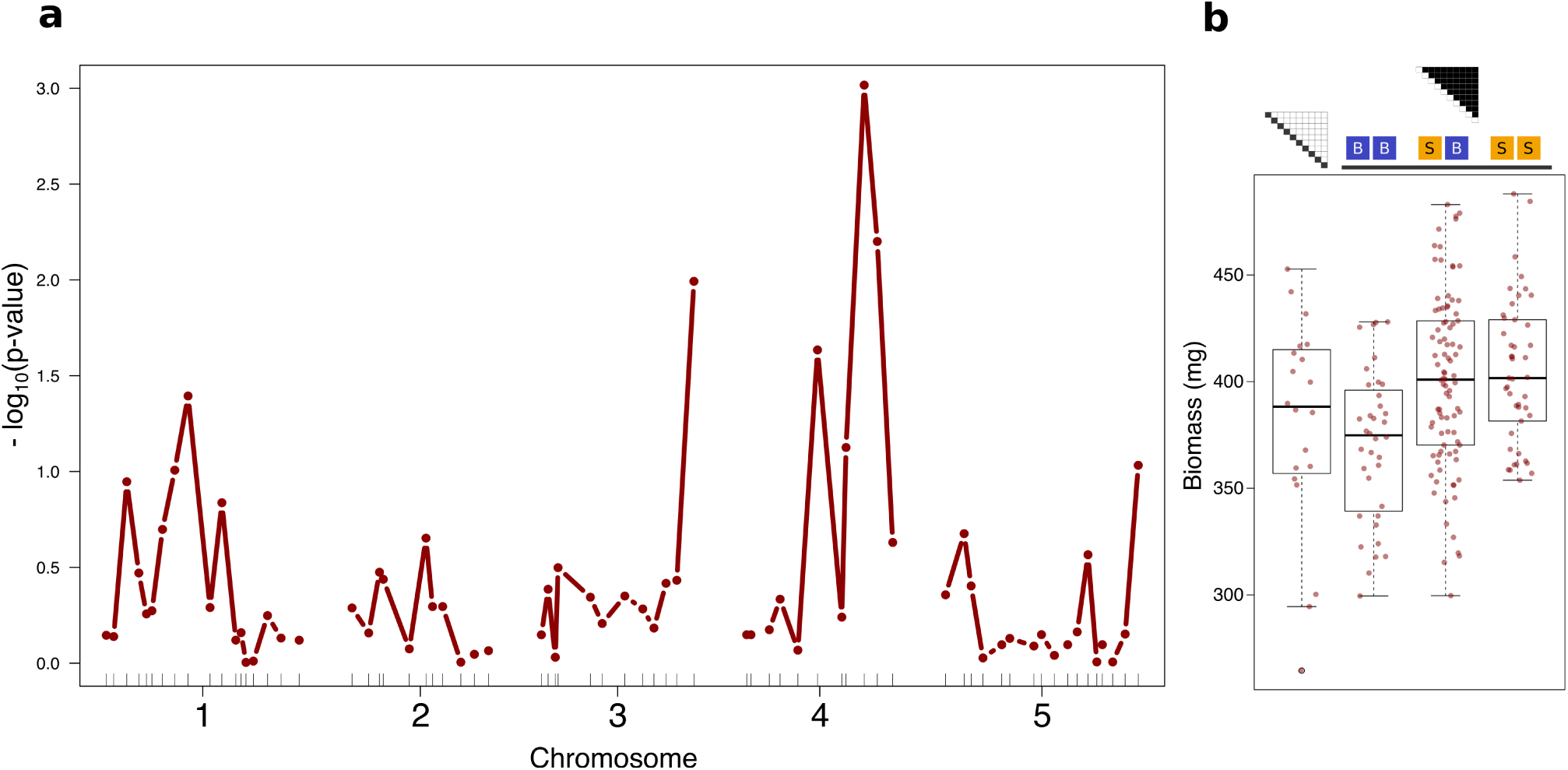
A major effect locus driving complementarity between genotypes. **a**, QTL mapping of SCA variation across allelic diversity levels using a marker regression technique by contrasting SCAs of mono-allelic RIL mixtures (BB and SS) with bi-allelic mixtures (BS). **b**, Pot-level productivity of each genotype composition (average of four blocks) in dependence of allelic composition at marker MSAT4.9. n_composition_ = 210

**Supplementary Figure 3.**
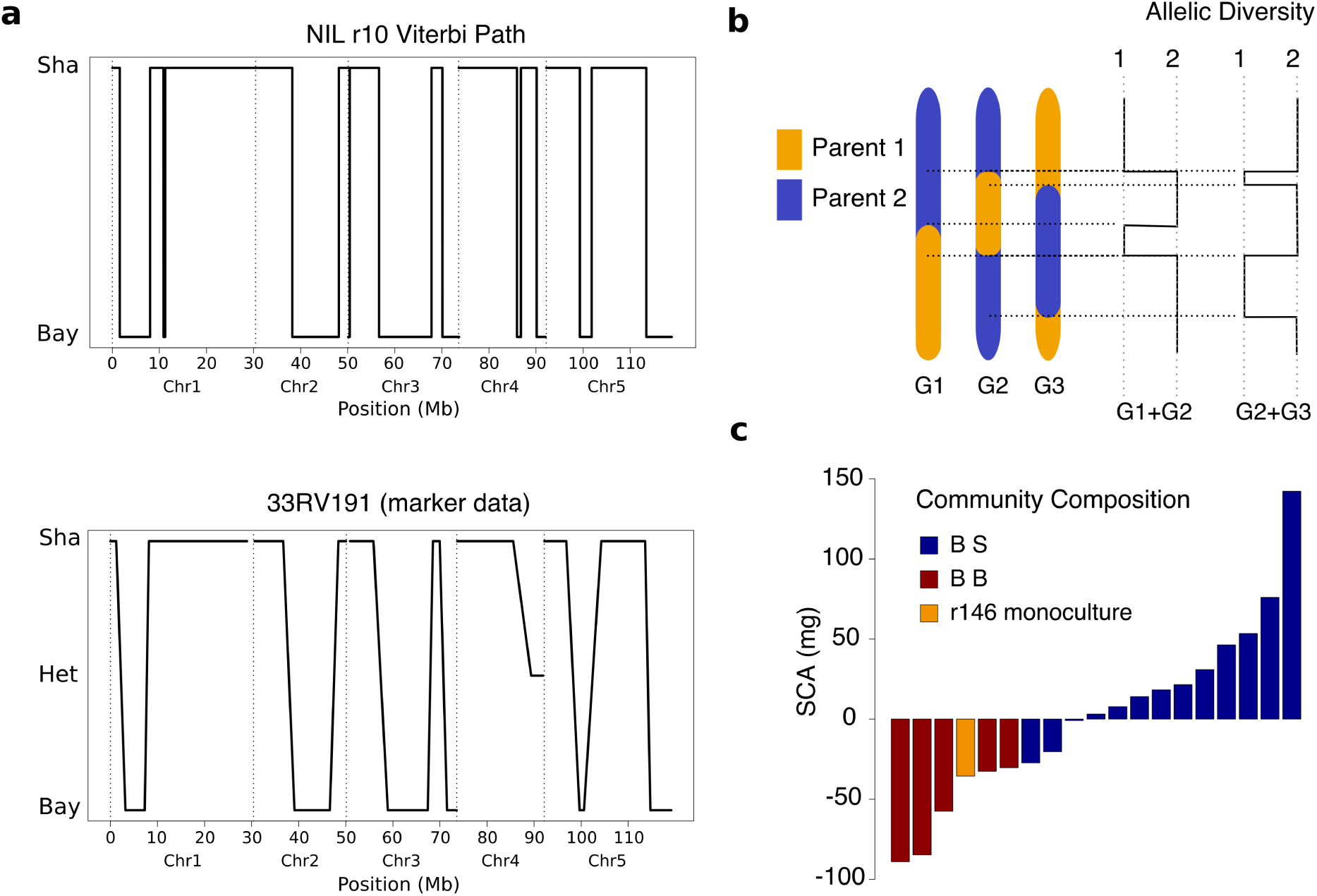
Fine-mapping and persistence of allelic diversity effects in near-isogenic line. **a**, The comparison of the reconstruced genotype of NIL r10 (HMM Viterbi-path across all chromosomes, homozygous on lower arm of chromosome 4) in comparison to publicly available molecular marker-based genotyping data of the ancestral line from which it was derived (heterozygous on lower arm of chromosome 4) – showing a high degree of congruence between the re-constructed genotype based on whole-genome resequencing and the marker data. **b**, Schematic outline of a possible cause of the high mapping resolution achieved through the diallel design. A major advantage of the design is the joint dependency of community-level allelic diversity on recombinations *within* and *between* recombinant inbred lines, such that mapping resolution increases very quickly. **c**, Extreme example of SCA variation across genotype mixtures all containing one specific genotype (NIL r146, homozygous for the Bay-allele at locus Chr4@16.92), either in combinations with NILs carrying the Sha-allele (dark blue bars) or the Bay-allele (dark red bars). Shown are the data from sand-rich soil only.

**Supplementary Figure 4.**
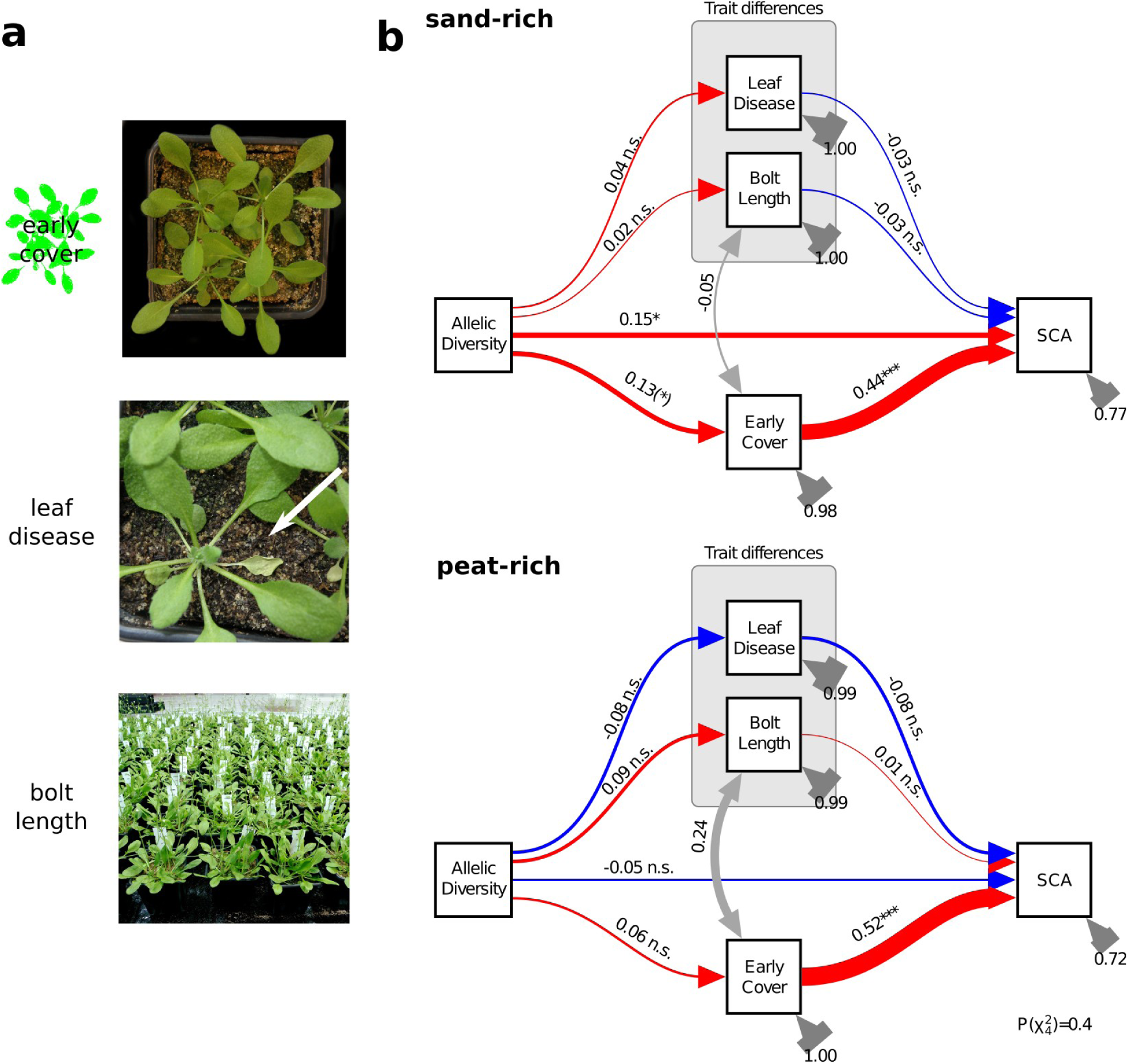
Observed above-ground trait variation does not explain overyielding of diallelic mixtures. **a**, Traits measured in the NIL diallel as proxy for productivity (early growth projected leaf cover), disease susceptibility (leaf disease at 30 days after sowing) or phenology (bolt length at 35 days after sowing) **b**, Path diagram of multi-group structural equation model showing direct effects of allelic diversity at locus Chr4@16.92 on SCA on sandy but not on peat soil. Red and blue arrows show positive and negative standardized path coefficients, respectively. n.s. = not significant. (*) = p<0.1; * = p<0.05; ** = p<0.01.

**Supplementary 5.**
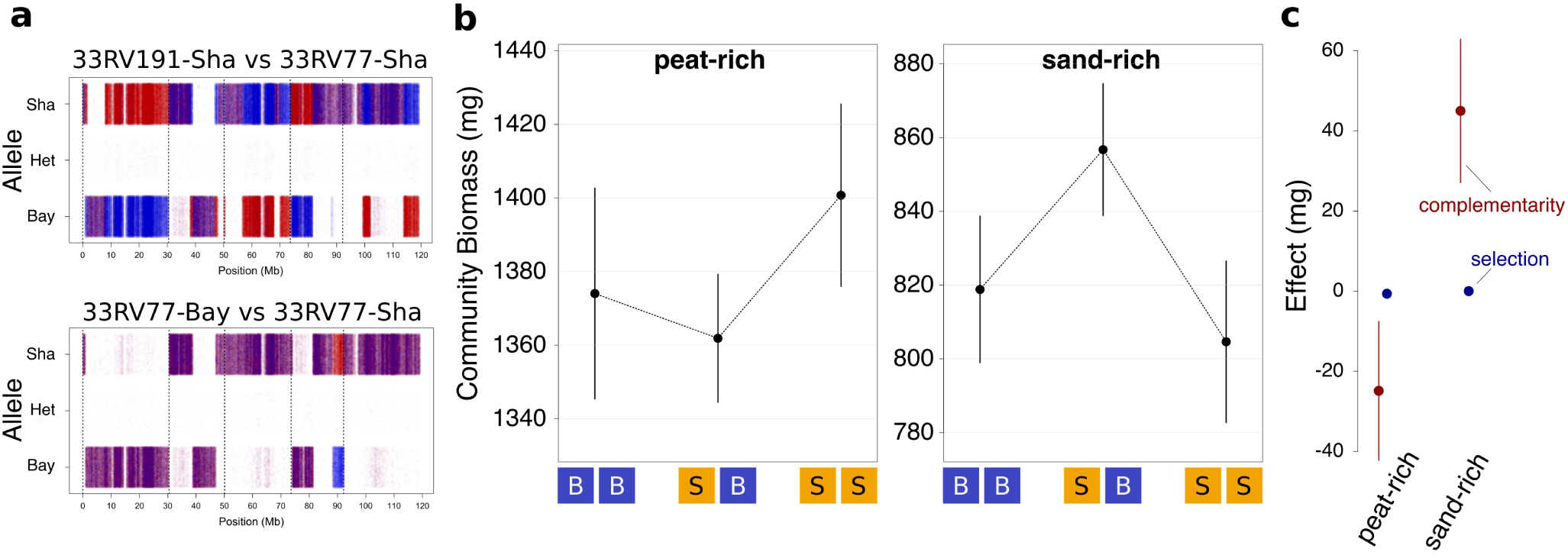
Soil-dependent effect of allelic diversity on overyielding and statistical complementarity and selection effects^S1^ in an independent near-isogenic background. **a**, Overlay of genotype calls at all polymorphic sites across the genome (Sha and Bay = homozygous for respective allele, het = heterozygous) of lines used for a mendelization. Top: a comparison of the genetically independent backgrounds 33RV191-Sha (in red, this background was used for the fine-mapping shown in Fig. 3) and line 33RV77-Sha (in blue) is shown. At the bottom, a comparison of the near-isogenic lines 33RV77-Sha (red) vs 33RV77-Bay (blue), the two lines that were used in the experiment shown in b and c. Purple regions depict overlap of genotype calls, vertical lines separate the different chromosomes. **b**, Mean ±s.e. of final aboveground biomass of communities consisting either of 33RV77-Sha plants only (BB), of mixed communities (BS), or of communities consisting of 33RV77-Bay plants only (SS) on either peat-rich (left) or sand-rich (right) soil. **c**, Complementarity and selection effects sensu Loreau and Hector calculated from the experiment shown in b.

**Supplementary 6.**
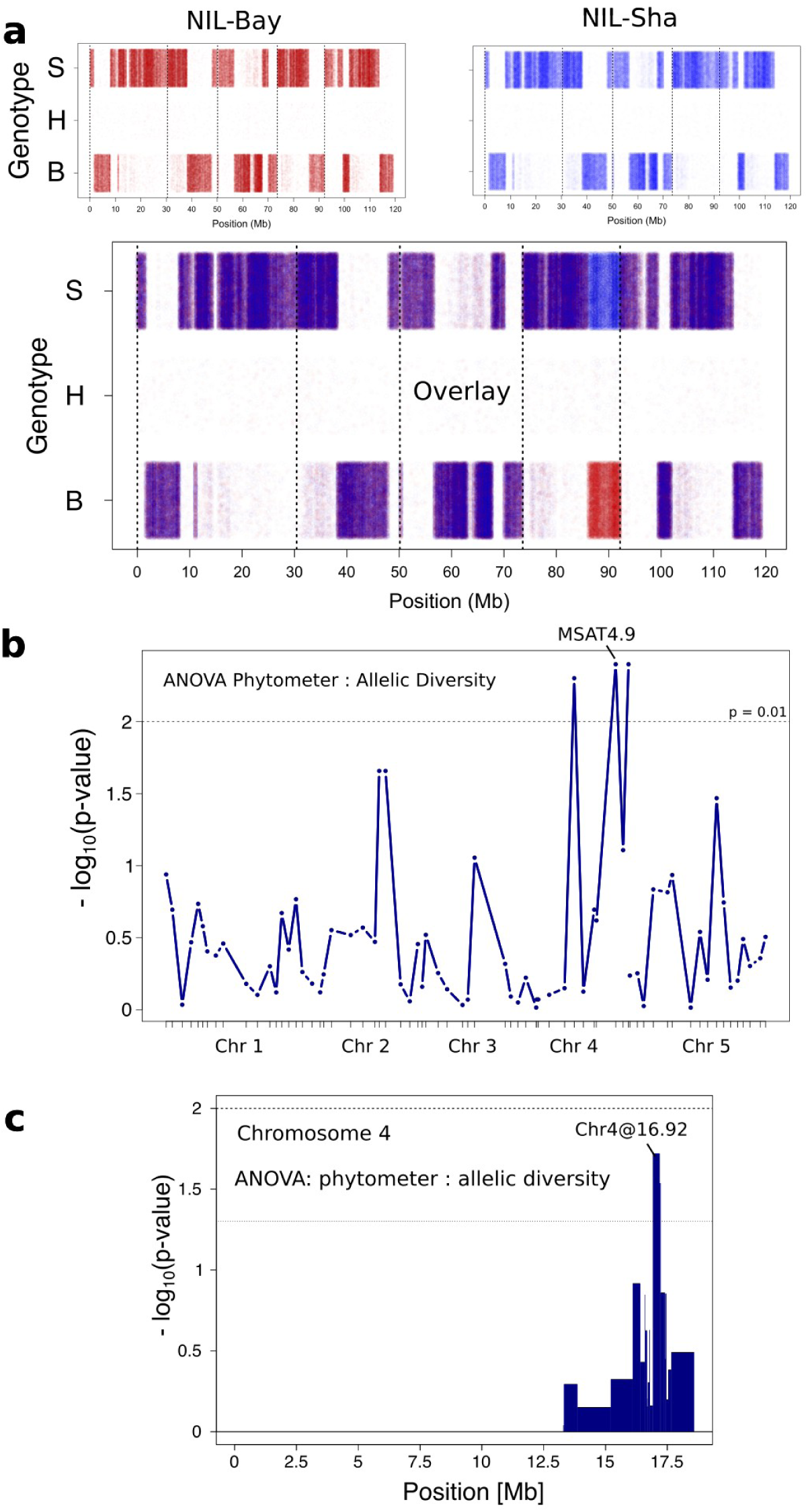
Soil feedback experiments. **a**, Genotype calls at all polymorphic sites across the genome (B = homozygous for the Bay allele, H = heterozygous, S = homozygous for the Sha allele) obtained by genome re-sequencing phytometer genotypes NIL-Bay (33RV191-Bay, top left, red) and NIL-Sha (33RV191-Sha, top right, blue). The overlay of the genotype calls of both lines (bottom) confirms that these lines are isogenic but for variation on lower arm of chromosome four. The two phytometers were employed on soil derived from the NIL diallel. **b, c**, QTL mapping by marker regression of phytometer-specific responses to soil legacy of previous generation allelic diversity (i.e. allele-diversity × phytometer [Phy×AD] interaction in Supplementary Table 1, but in a model without adjusting for diallel pot biomass). The mapping of diversity effects through such influences on soil legacy could have been applied to the identification and fine-mapping of the same major effect locus, without ever measuring biomass productivity (albeit with slightly relaxed statistical criteria). Shown are negative log_10_-transformed P-values for each marker position in the RIL (b) or genotype block in the NIL (c) diallels.

## Supplementary Discussion

As outlined in Figures 1 and 4, and as described in the Methods, we performed two soil-feedback experiments to test whether allelic diversity effects extend across generations. We accounted for potential effects explainable by variation in plant productivity during the soil training phase (e.g. nutrient draw-down or environmental correlations) in linear models (terms *diallel biomass* and *diallel biomass*×*soil* as described in the Methods section). Significantly different soil conditioning through allelic diversity, as assessed by phytometer performance in a next growing period, was interpreted as an “extended phenotype”^S2^ (i.e. a legacy of allelic mixture that persists through time in the soil, even after removal of the original plant communities). These soil factor-mediated allelic legacy effects interact with phytometer genotype, giving rise to phytometer-specific responses (term “Phy×AD” in Supplementary Table 1).

As we emphasize, the specific mechanisms underlying both soil training and phytometer responses await further experiments, since the response patterns (Fig. 4) do not allow for a simple mechanistic model. One reason why it is difficult to infer specific mechanisms (e.g. specific soil factors, and their interactions with genetic variants) is the number of variables that differed between the two experiments because of constraints on experimental design: 1) environmental conditions (different calendar dates, different batches of soil); 2) genetic variation in the two diallel population (RILs vs NILs), potentially resulting in different effects of epistasis; and 3) phytometer genotypes (parental lines vs NILs).

In the RIL diallel, we used the two parental accession Bay and Sha as phytometers and a split-plot design to test for differential responses of these phytometer to soil conditioning. After the NIL diallel, we used two near-isogenic phytometer (33RV191-Bay and 33RV191-Sha) as phytometers in a similar test for differential responses to soil conditioning. Naively, one might expect approximately congruent responses of the phytometers carrying the same alleles at the diversity locus under question, i.e. the parental genotype Bay in the RIL diallel (Figure 4 b, yellow lines) and the NIL-Bay genotype in the NIL diallel (Figure 4 c, yellow lines) might respond similarly to the legacy of allelic diversity. However, this was not the case. For example, the Bay genotype grown on RIL diallel soil responded positively to conditioning by allelic diversity, whereas the NIL-Bay genotype in the NIL diallel responded somewhat negatively to conditioning by allelic diversity. It is noteworthy that the pattern found for Bay on RIL diallel soil is opposite to what would be expected if soil legacy was driven by a simple productivity-related depletion of resources in the conditioning phase; this suggests that allelic diversity alleviates a negative soil factor (e.g. inhibits enemies), or promotes a positive factor (e.g. mutualistic organisms).

Despite differences, the RIL and NIL soil-feedback experiments nevertheless convincingly demonstrate i) that allelic diversity effects extend across time through soil conditioning, and ii) that phytometer genotype determines the response to such conditioning (phytometer×allelic diversity interaction, both experiments, see Table S1). The fact that allelic diversity effects within a community and allele-specific legacy effects on individuals separated in time map to the same locus suggests to us that both are related to similar mechanisms.

